# An integrated approach to protein discovery and detection from complex biofluids

**DOI:** 10.1101/2022.01.03.474834

**Authors:** Gordon T. Luu, Chang Ge, Yisha Tang, Kailiang Li, Stephanie M. Cologna, Joanna E. Burdette, Judith Su, Laura M. Sanchez

## Abstract

Ovarian cancer, a leading cause of cancer related deaths among women, has been notoriously difficult to routinely screen for and diagnose early. Researchers and clinicians continue to seek routinely usable, non-invasive, screening methods as early detection significantly improves survival. Biomarker screening is ideal; however, currently available ovarian cancer biomarkers lack desirable sensitivity and specificity. Furthermore, the most fatal forms, high grade serous cancers often originate in the fallopian tube; therefore, sampling from the vaginal environment provides more proximal sources for tumor detection. To address these shortcomings and leverage proximal sampling, we developed an untargeted mass spectrometry microprotein profiling method and identified a signature of cystatin A, validated this protein in an animal model, and sought to overcome the limits of detection inherent to mass spectrometry by demonstrating that cystatin A is present at 100 pM concentrations using a label-free microtoroid resonator. The findings highlight the potential utility for early-stage detection where cystatin A levels would be low.

**Significance Statement:** It is now clear that high-grade serous ovarian cancer can originate in the fallopian tube epithelium. These tumors colonize the ovary and then metastasize throughout the peritoneum. This discovery has raised important, and yet unaddressed, questions how we might be able to detect and screen for this deadly disease for which there is no routine screening. We have leveraged vaginal lavages from a murine model of the disease as a complex biological fluid for untargeted discovery of microproteins using mass. We improved our limits of detection by conjugating a cystatin A antibody to the surface of a microtoroid resonator to allow us to specifically detect cystatin A from vaginal lavages at early time points across biological replicates.

## Introduction

Ovarian cancer remains one of the most lethal gynecological cancers in women, with 313,959 new cases worldwide in 2020; it is second only to cancer of the cervix uteri, which had 604,127 new cases in 2020^1^. Due to a paucity of early detection strategies, the five-year relative survival rate from 2011-2017 for ovarian cancer (49.1%) was nearly half that of uterine cancer (81.1%)^2–4^. When detected early, the chance of survival for ovarian cancer can increase to almost 90%^5^. Ovarian cancer seldom displays clinical symptoms prior to metastasis, leaving patients unaware of their condition until the disease progresses to stage III or IV. However, currently available screening methods are invasive and costly (i.e. transvaginal ultrasounds, biopsies) or lack the specificity and sensitivity to be used routinely (i.e. cancer antigen 125, human epididymis protein 4, OVA1®, Overa®)^6–9^. Therefore, there is an urgent need for 1) reliable biomarkers accompanied by methodologies to detect them and 2) novel sampling methods to routinely screen for early-stage gynecological cancers.

Brinton *et al*. suggested that local tumor microenvironments may be a more appropriate alternative sampling sources for ovarian cancer biomarkers as they would prove to be more useful in the detection of primary tumors, which cannot be done using metastatic biomarkers^10^. To that end, Costas *et al*. have previously suggested novel sampling methods in the context of endometrial cancer and outlined several criteria for effective sampling, including 1) high-throughput, 2) able to detect early-stages of disease, 3) minimally invasive, and 4) affordable^11^. Vaginal sampling may provide a site where tumors that arise in the fallopian tube (i.e. high grade serous ovarian cancer, a prominent and particularly fatal subtype of ovarian cancer; HGSOC) or in the uterus (i.e. endometrial cancer) may escape and concentrate earlier than in serum^12–14^.

We have previously used vaginal lavages to collect intact cells and extracellular proteins from the vaginal microenvironment of mice with OVCAR-8-RFP tumors, which models HGSOC (**Figure 1a**). Vaginal lavages collected over eight weeks from a cohort of five mice were analyzed using matrix-assisted laser desorption/ionization time-of-flight mass spectrometry (MALDI-TOF MS) to profile the microprotein range, which consists of proteins < 30 kDa. Here, earlier time points (Days 0-28 post tumor implantation) and later time points (Days 35-56 post tumor implantation) represented early-stage and late-stage ovarian cancer, respectively. Differentially expressed microproteins were present and differentially expressed during tumor progression. However, a limitation of MALDI-TOF MS protein profiling is that it only provides a putative intact mass with no other information on protein identity^6^. Herein, we have used liquid chromatography-tandem mass spectrometry (LC-MS/MS) based bottom-up proteomics to analyze the same murine lavages reported by Galey *et al*. to identify these putative biomarker features (**Figure 1b**). In doing so, we have identified cystatin A and confirmed its presence in tumors and murine reproductive tissue which provided insight into its spatial distribution *in vivo*. Knowing that MALDI-TOF MS provides a limited means for early detection for specific biomarkers over the multidimensional fingerprinting, we sought to integrate more sensitive, specific technologies for detection from the same lavage samples. Therefore, we applied a targeted approach to leverage frequency-locked optical whispering evanescent resonator (FLOWER) as a potential alternative early-stage screening method (**Figure 1b**)^15–17^. FLOWER is based on microtoroid optical resonator technology and allows for label-free detection of potentially attomolar concentrations of an analyte, which is not possible with ELISA or MS (**Figure 1c-e**)^17–19^. With FLOWER, light is evanescently coupled into the microtoroid using a tapered optical fiber. At the resonance frequency of the microtoroid, there will be a dip in the transmission of the light that passes through the optical fiber. This transmission dip is monitored as analyte molecules bind to antibodies anchored to the surface of the toroid. One advantage of using FLOWER for these experiments is its ultra-sensitivity and compatibility with small (μL) amounts of biofluids (i.e. murine vaginal lavages)^18–20^. In sum, we present a workflow for combining powerful untargeted technologies to discover microproteins for use with specific, sensitive label free targeted screening.

**Figure 1.**
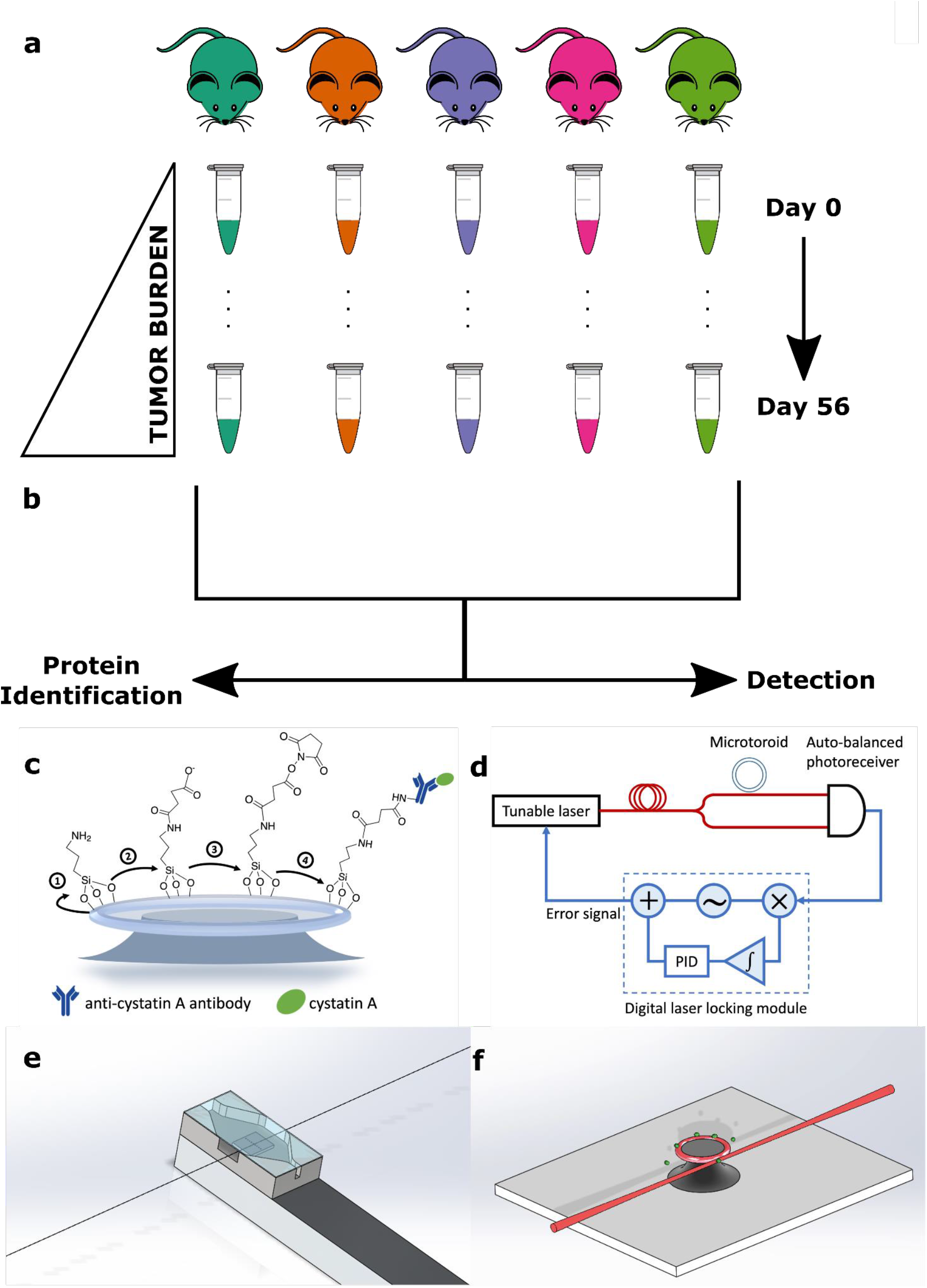
(a) Previous study by Galey et al. collected murine vaginal lavages for xenograft OVCAR-8-RFP mice over eight weeks and profiled each lavage using MALDI-TOF MS. (b) The current study seeks to identify proteins detected from MALDI-TOF MS protein profiles and use more sensitive detection techniques. (c-f) Schematic of cystatin A bio-sensing experimental setup using FLOWER. (c) Preparation of the microtoroid for binding to the cystatin A - Ab complex. (d) Block diagram of the FLOWER system. A digital laser locking module enables adaptive tracking of the microtoroid’s resonance frequency which changes as analytes bind to its surface. (e) Schematic representation of the microfluidic chamber designed to allow the passage of the optical fiber. (f) An isometric view showing analyte molecules (green) binding to the surface of the toroid. The toroid is shown positioned next to a tapered optical fiber for evanescent coupling of light into the toroid.

## Results

### Annotation of Differentially Expressed Microproteins in Murine Vaginal Lavages via LC-MS/MS Based Bottom-Up Proteomics

To annotate the microproteins from previously collected murine vaginal lavages, LC-MS/MS based bottom-up proteomics was used with the goal of identifying the putative proteins detected via MALDI-TOF MS screening. It should be noted that a typical murine vaginal lavage has a volume less than 200 μL with variable amounts of biological material (i.e. cells, proteins) across biological replicates. Due to the limited sample volumes alternating timepoints (Days 7, 21, 35, and 49) were pooled (N=3) for proteomics experiments. Limited sample digestion was performed with a modified filter aided sample preparation (FASP) protocol with Lys-C as the digestion enzyme. In solution enzymatic digestion for pooled Days 35 and 49 samples was also performed as an alternative enzymatic digestion method. LC-MS/MS followed by MaxQuant (version 1.5.4.0) to search the resulting peptide sequences against UniProt proteomes for *Homo sapien, Mus musculus*, and *Rattus norvegicus* (**Table 1**).

**Table 1.** Curated list of proteins annotated from LC-MS/MS data processed via MaxQuant. Proteins < 2 unique peptides and > 32 kDa were filtered out. Six proteins found in MALDI-TOF MS protein profiles are shaded in green. Q-values for all annotations (not shown) were found to be 0.

| Protein | Gene | Molecular Weight [kDa] | Sequence Coverage[%] | Unique Sequence Coverage [%] | Sequence Length | Score |
| --- | --- | --- | --- | --- | --- | --- |
| Annexin | Anxa2 | 31.263 | 12.9 | 4.4 | 272 | 2.5 |
| Caspase-14 | CASP14 | 27.679 | 14.9 | 14.9 | 242 | 7.4 |
| Histone H1.2/H1.3/H1.4 | H1-2/H1-3/H1-4 | 21.316 | 16.0 | 16.0 | 212 | 3.5 |
| Calmodulin-like protein 5 | CALML5 | 15.892 | 47.3 | 47.3 | 146 | 60.3 |
| Gp dh N domain-containing protein/Glyceraldehyde-3-phosphate dehydrogenase | Gadph | 15.373 | 5.6 | 5.6 | 142 | 2.9 |
| Fatty acid-binding protein 5 | FABP5 | 15.164 | 24.4 | 24.4 | 135 | 9.7 |
| Protein S100-A9 | S100A9 | 13.242 | 24.6 | 24.6 | 114 | 28.1 |
| Histone H2B |  | 13.134 | 16.9 | 16.9 | 118 | 43.3 |
| Protein S100-A7 | S100A7 | 11.471 | 28.7 | 28.7 | 101 | 6.3 |
| Histone H4 | H4c1 | 11.367 | 26.2 | 26.2 | 103 | 2.0 |
| Dermicidin | DCD | 11.284 | 16.4 | 16.4 | 110 | 5.5 |
| Cystatin-A | CSTA | 11.006 | 39.8 | 39.8 | 98 | 16.2 |
| Protein<br>S100-A8 | S100A8 | 10.834 | 45.2 | 45.2 | 93 | 17.3 |
| Ubiquitin-60S<br>ribosomal<br>protein L40 |  | 7.132 | 49.2 | 49.2 | 63 | 111.6 |

### Retrospective Analysis of MALDI-TOF Protein Profiles Revealed Microproteins of Interest

Manual retrospective analysis was conducted due to the relatively small list of annotated proteins. The dataset reported by Galey *et al*. was reprocessed using MALDIquant to generate a feature matrix, which was found to contain six of the annotated proteins from LC-MS/MS experiments (highlighted in **Table 1**)^20^. Of these six proteins, two in particular proved to be of interest based on previously reported biological significance in the literature for ovarian cancer and other types of cancer: protein S100-A8 and cystatin A. Protein S100-A8 (*m/z* 10835 +/− 40 ppm, **Figure S1**) was found to be downregulated over time in vaginal lavages as tumor burden increased, while cystatin A (*m/z* 11007 +/− 30 ppm) was found to be upregulated over time. Cystatin A was chosen for further analysis and experiments since a protein that is upregulated in the vaginal microenvironment is a preferable biomarker. **Figure S2** shows longitudinal tumor burden as measured by *In Vivo* Imaging System (IVIS) and intensity of cystatin A as measured by MALDI-TOF MS. Interestingly, tumor burden in Mouse 902 appeared to plateau and slightly decrease after Day 42. A similar trend was observed in cystatin A intensity for Mouse 902 as detected by MALDI-TOF profiling (**Figure 2a** and **2b**). Although Mouse 903 showed no decrease in tumor burden, its cystatin A intensity showed a similar decrease in intensity to that of Mouse 902.

**Figure 2.**
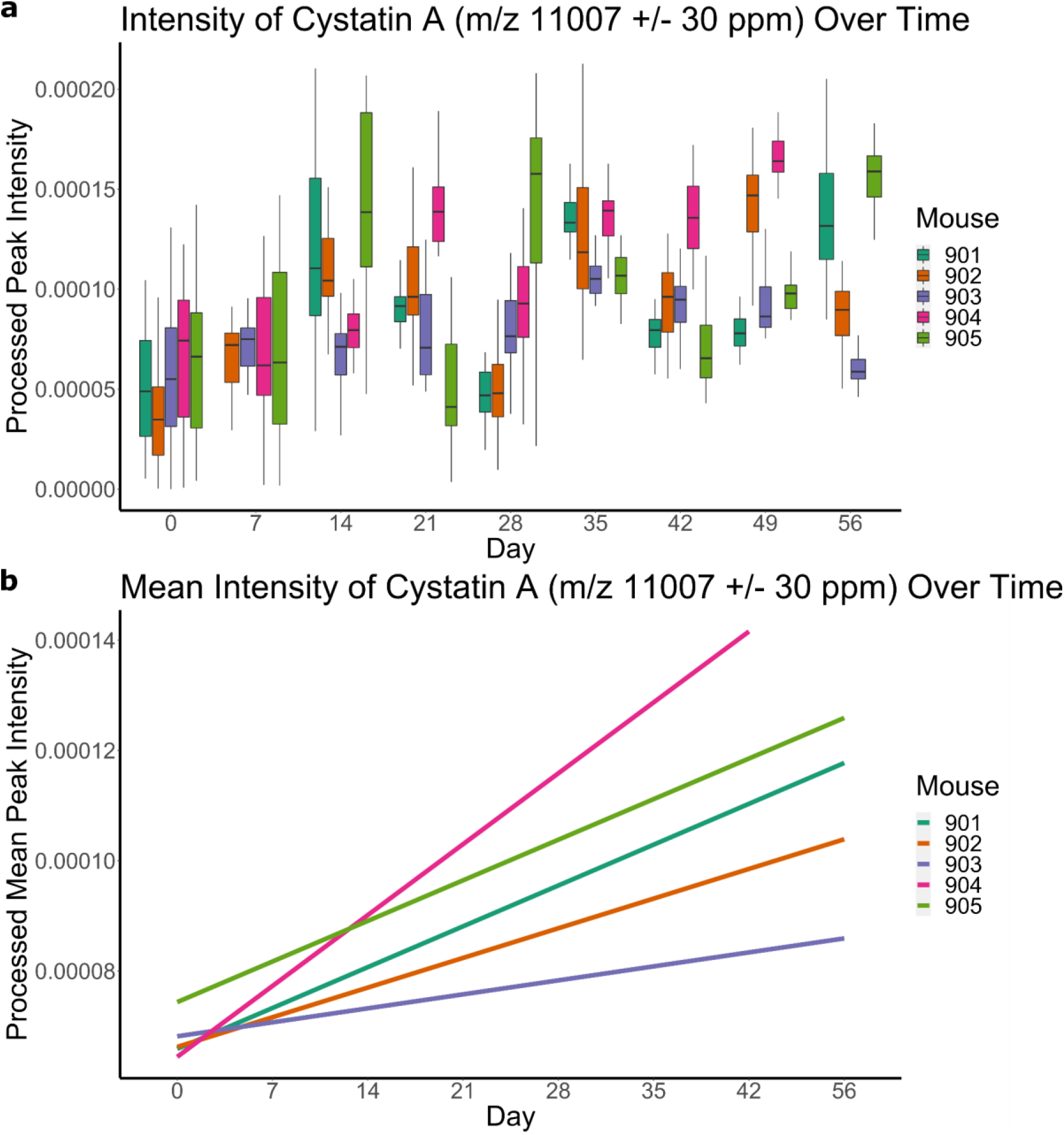
Feature intensity as measured by MALDI-TOF MS of *m/z* 11007 (cystatin A) in protein profiles in five biological replicate xenograft murine models transfected with OVCAR-8-RFP tumors. (a) Box plot showing the intensity of cystatin A at each corresponding time point. (b) Linear regression trend lines for the mean intensities of cystatin A at each time point showing upregulation.

### Immunohistochemistry Staining Reveals Presence of Cystatin A in Murine Reproductive Tissue and Xenografted Tumors

To validate the presence of cystatin A *in vivo*, tumors and reproductive tract tissue were stained for IHC with antibodies directed against cystatin A (**Figure 3a**). All of the tumors showed varying levels of cystatin A protein expression at the edges of the tumor with a unique “spotting” pattern of distribution; interestingly, this spotting, and therefore cystatin A, was found throughout the tumor collected from Mouse 903. Overall, the presence of cystatin A was confirmed in all the tumors but its localization and abundance lacked consistency. Importantly, OVCAR8 cells are RFP tagged and no RFP was detected in the lavages indicating that proteins in the lavage are reflective of the tumor microenvironment rather than the tumor cells.

**Figure 3.**
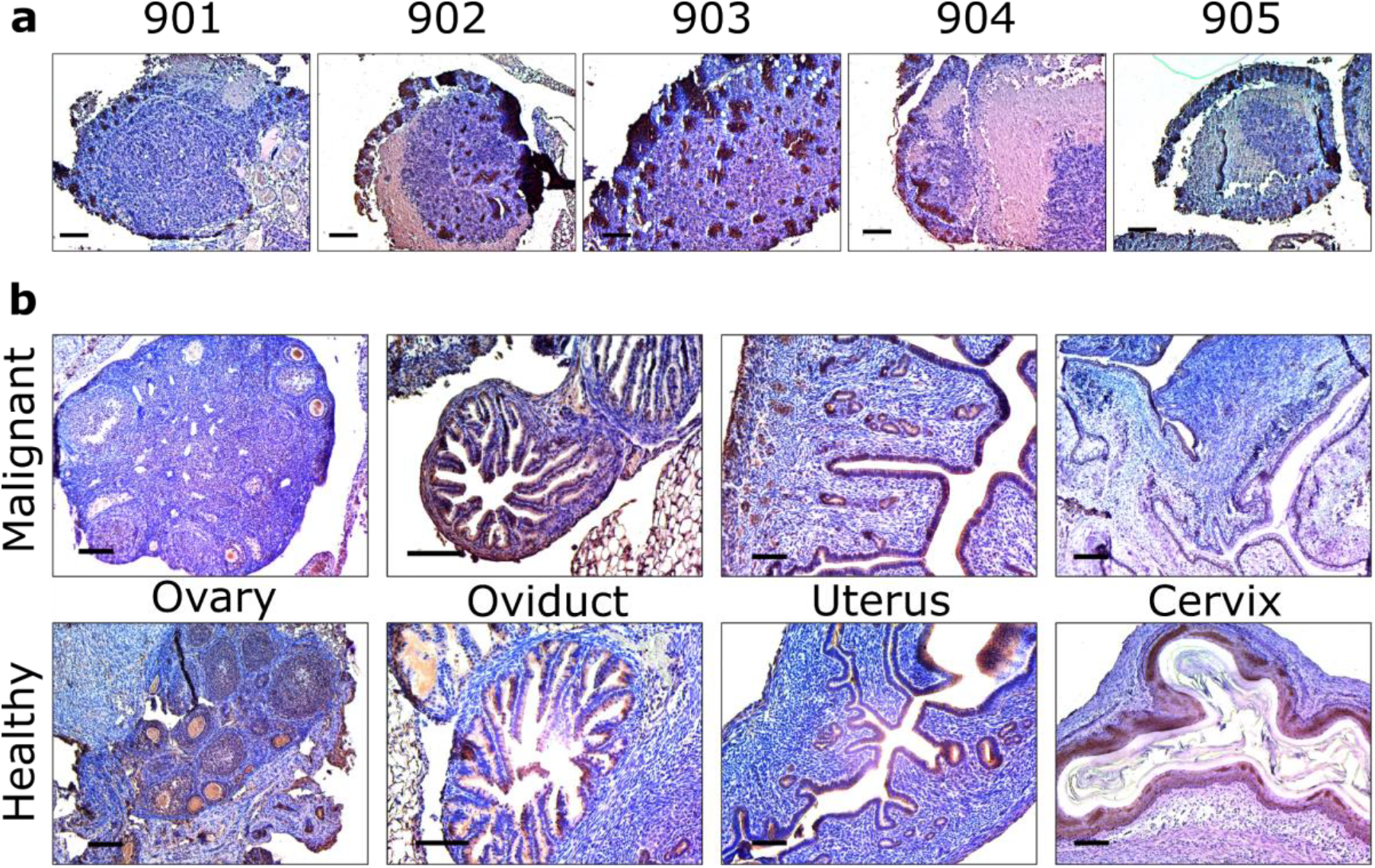
IHC staining for (a) tumors from Mouse 901-905 and (b) healthy and malignant murine reproductive tissue (ovary, oviduct, uterus, and cervix).

Similarly, IHC staining was also performed on reproductive tissue taken from healthy mice and those with OVCAR8 tumors (ovary, oviduct, uterus, and cervix; **Figure 3b**). Here, cystatin A can be found in the epithelial cells of the fimbriae of the oviduct and in tissue lining the lumen of the uterus. In oviducts, where OVCAR8 tumors were present, cystatin A abundance was slightly increased, and in the malignant uterus, cystatin A increases along the edge of the tissue and the spotting pattern seen in tumors from **Figure 3a** is also present. In the healthy ovary, cystatin A can be found around the tissue edges and “pooling” near the center of the tissue section. Lastly, the healthy cervix shows large amounts lining epithelial tissue that is absent in the malignant cervix. Although the ovaries do not appear to have much cystatin A, differential expression is observed in the other types of reproductive tissue.

### FLOWER Allows for Detection of Low Concentrations of Cystatin A in Limited Biological Samples

FLOWER was also used as an orthogonal method to detect cystatin A. The binding curves for all murine vaginal lavages from Days 14, 28, 42, 56, and a 100 pM cystatin A reference standard were recorded (**Figure S3**). In Mouse 905, there is an increase in cystatin A from Day 14 to Day 28. Additionally, compared to the 100 pM standard, later time points (Days 28 to 56) display equal or greater initial slope values, which proves to be helpful in the quantification and estimation of cystatin A levels and concentrations. To characterize cystatin A levels in each sample, the binding curves were fit with either a linear function

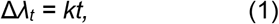

where *k* is the initial slope and *t* is time, or an exponential growth equation

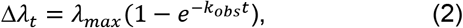

where Δ*λ_t_* is the time-dependent resonance wavelength shift and *λ_max_* is the maximum wavelength shift at the plateau (**Figure S4**). The term *k_obs_* is the observed rate constant for association^21^. To obtain the binding rate, the exponential function was differentiated at time t = 0, which can be expressed as the initial slope *k* (*k* = *λ_max_ x k_obs_*).

To minimize the effect of any variations due to 1) the size, geometry, and characteristics of different microtoroids, 2) flow conditions of the sample, and 3) the immobilization efficiency, which potentially influences the combined dataset, we calibrate all the initial slope values by normalizing the values obtained from the response from a standard 100 pM injection in each sensing trace^22^. Here, the initial slope values obtained from the scatter plot and linear regression lines in **Figure 4** are plotted as a function of days since tumor implantation for all five mice. With the exception of Mouse 903, the initial slope values over time show a progressive growth, revealing an overall increase in cystatin A levels (**Figure 4a**). Mouse 903, however, showed low levels of cystatin A from Day 0 to Day 42, followed by a sudden increase in the initial slope from −5.676 to 25.157 fm/s at Day 56, which is consistent with MS data. Low concentrations of cystatin A in Mouse 903 is most likely attributed to biological variance and/or sampling error. Using the 100 pM cystatin A standard’s initial slope as a reference, cystatin A levels in the diluted vaginal lavages of the remaining four mice are greater than or equal to 100 pM at 42 days post xenograft. Since murine vaginal lavages were diluted a thousand-fold, estimated cystatin A levels are actually greater than or equal to 100 nM at Day 42.

**Figure 4.**
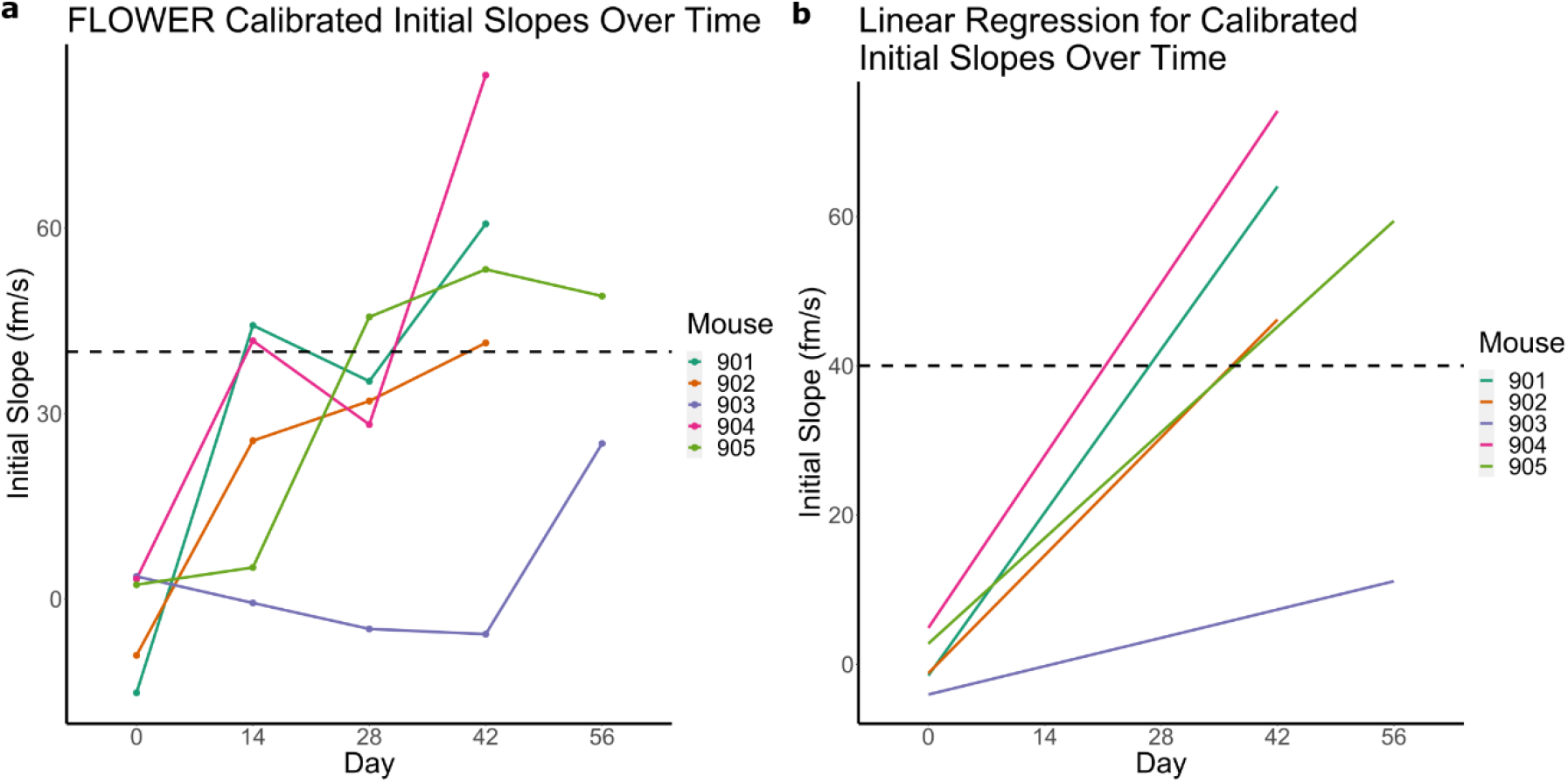
Label-free detection of cystatin A using FLOWER. The combined plot for all five mice samples with 100 pM calibration. (a) The scatter plot of the initial slope values as a function of days since tumor implantation. The dashed line indicates the value obtained from the normalized standard concentration of 100 pM. (b) Linearly fitted initial slope values versus time.

## Discussion

MALDI-TOF MS protein profiling is a powerful tool that is leveraged in a variety of clinical and academic settings using instruments such as the Biotyper (Bruker Daltonics) and VITEK (bioMerieux)^5^. FDA approved protocols have been in use and allow for high confidence identification of bacteria and yeast^23^. Similarly, the Mass-Fix assay has been developed and used clinically to monitor monoclonal proteins (M-proteins) in serum collected from plasma cell dyscrasias patients using MALDI-TOF MS protein profiling^24–28^. Multiple myeloma, a cancer that forms from plasma cells, is a prevalent plasma cell dyscrasia that also has a low five-year survival rate (55.6%)^29,30^. By analyzing serum, which is devoid of red blood cells, white blood cells, erythrocytes, and clotting factors, Mass-Fix monitors a relatively simple biofluid. Petukhova *et al*. has previously optimized a method for the detection of microproteins from a more complex biofluid (i.e. whole cell cultures) in which a mixture of cells are present and analyzed using MALDI-TOF protein profiling^12^. A follow up study by Galey *et al*. used MALDI-TOF MS protein profiling to identify features that are differentially expressed over time as tumor burden increased in murine vaginal lavages^11^. Here, protein profiling was shown to be a viable method for detecting statistically significant features of interest even with increased sample complexity.

In the present study, we have detected and annotated cystatin A (also known as stefin A or acid cysteine proteinase inhibitor; ACPI) among other microproteins from murine vaginal lavages containing cells and biomolecules from the vaginal microenvironment. Although cells were present, we have previously shown that fluorescently labeled tumor cells were not detected via microscopy or flow cytometry despite their close proximity to the sampling site, indicating the differential expression of microproteins over time were likely the result of changes in the vaginal microenvironment and not the OVCAR-8 tumors themselves^11^. Cystatin A is a human cysteine proteinase inhibitor first identified from the cytosol of human polymorphonuclear granulocytes and has been shown to play roles in various cancers (i.e. colorectal, breast, non-small-cell lung) and diseases (i.e. psoriasis, glaucoma)^31–43^. As a type 1 cystatin, cystatin A is normally found intracellularly; however, it has also been shown to appear in biofluids^44,45^. MALDI-TOF MS protein profiling revealed differential expression of cystatin A, which was annotated using LC-MS/MS based bottom-up proteomics. Lah *et al*. have previously observed upregulation of cystatin A in ascites fluid (buildup of excess fluid in the abdomen), while Kastelic *et al*. have observed downregulation of cystatin A in ovarian carcinoma (derived from epithelial tissue)^46,47^. Therefore, the increase of cystatin A in the vaginal microenvironment over time was consistent with data reported by Lah *et al*. as we have also observed upregulation of cystatin A in fluids derived from the vaginal microenvironment (i.e. vaginal lavages). Additionally, the downregulation of cystatin A observed in malignant cervical epithelial tissue is consistent with Kastelic et al. IHC staining was used to validate the presence of cystatin A in tumors, healthy, and malignant murine reproductive tissue; furthermore, cystatin A appeared to be localized to epithelial cells lining most of the reproductive tract.

After observing upregulation of cystatin A in protein profiles and validating its presence in tissues, we employed FLOWER as an orthogonal method of detection to explore whether we could measure cystatin A more reliably from earlier time points and determine the limit of detection. We found that FLOWER detected the presence of cystatin A at pM concentrations. Early stages of disease are often difficult to detect due to our inability to reliably detect changes in protein and metabolite expression downstream of genetic mutations with current methods (i.e. MALDI-TOF MS, enzyme-linked immunosorbent assay; ELISA). In diseases such as ovarian cancer, early screening of disease can be correlated to improved patient prognosis^48^. Therefore, the development of label-free technologies capable of lower limits of detection for specific analytes are important towards the goal of screening for early stage disease.

Based on the data presented here, the combination of untargeted (MALDI-TOF MS protein profiling) and targeted (FLOWER) detection methods provides a power platform that has led to the detection of cystatin A, a promising candidate ovarian cancer biomarker. As we continue to discover novel biomarkers, they can be used in conjunction with or as a replacement for known biomarkers in multimarker panels. Furthermore, these detection methods can be multiplexed with other current (i.e. biomarker panels) or yet to be developed technologies to further improve our ability to screen for disease in a clinical setting, which opens the potential for larger clinical cohorts and allows for the ability to perform retrospective studies to aid in biomarker discovery. With that being said, the limitations of the current study must also be considered. Due to the limited nature of the vaginal lavages, sample pooling was required for sufficient material to perform bottom up proteomics experiments. As observed by Diz *et al*., pooling samples can provide benefits including reduction of biological variance in biological replicates, but it also results in reduced statistical power and can leave low abundance proteins undetected^49^. Another limitation of the current study is the lack of tissue to correlate with every lavage time point. Lastly, targeted detection via FLOWER requires the presence of recombinant monoclonal antibodies specific to the protein of interest on the microtoroid, which may prove difficult for proteins without commercially available antibodies. Nevertheless, a major advantage of our study was the lavages and tissues were all collected from the same mice used previously for fingerprinting. We leveraged the limited biological fluids for both identification, validation, and specific, sensitive detection which also highlighted the biological variability of this disease in our model organisms. We believe the reported workflow has a high potential to achieve a routine screening method for diseases such as ovarian cancer.

## Materials and Methods

### Chemicals

Aminopropyl trimethoxysilane (APTES, 440140), acetic acid (695092), succinic anhydride (239690), dimethyl sulfoxide (DMSO, 276855), anhydrous ethanol (676829), ethanolamine (E6133) were purchased from Sigma Aldrich. Pierce premium grade N-hydroxysulfosuccinimide (sulfo-NHS, PG82071) and 1-ethyl-3-(3-dimethylaminopropyl) carbodiimide (EDC, 22980) were purchased from Thermo Fisher Scientific.

### In Vivo Murine Xenograft Study

Full details for the *in vivo* murine xenograft study have been previously reported11. During the study, vaginal lavages for each mouse (numbered 901-905) were collected at timepoints every seven days starting on Day 0 and ending on Day 56.

### Storage of Murine Vaginal Lavages

Murine vaginal lavages for mice 901 and 902 were normalized to 10,000 cells/μL, while mice 903, 904, and 905 were normalized to 5,000 cells/μL; the total volume of samples ranged from 4 μL to 20 μL, with one sample being 1 μL. All lavages were stored at −80° C until used. Day 7, 21, 35, and 49 were used for LC-MS/MS based bottom-up proteomics, while Day 14, 28, 42, and 56 lavages were shipped and used for detection of cystatin A via FLOWER. Upon receiving, the murine vaginal lavages were stored at −80° C.

### Preparation of Vaginal Lavages for Enzymatic Digestion

Protein concentration for samples were determined using a bicinchoninic acid (BCA) assay. Due to the low individual concentrations of each replicate, three biological replicates were pooled by timepoint (days). Pooled samples were diluted to a total volume of 500 μL and transferred to Amicon Ultra-0.5 30 kDa centrifugal filter devices (Millipore Sigma). Filter devices were centrifuged at 14000 g for 10 min. The process was repeated with a 100 μL and two 50 μL PBS washes under the same centrifugation conditions to ensure adequate sample collection. The retentate was also collected by inverting the Amicon spin column filter and centrifuging at 1000 g for 2 min. The retentate and filtrate were collected into clean 2 mL microcentrifuge tubes and dried *in vacuo* at 45° C for 1 hr.

### Enzymatic Digestion of Murine Vaginal Lavages Using S-Trap

A modified filter aided sample preparation protocol using the S-Trap (ProtiFi) was used for enzymatic digestion of all pooled murine vaginal lavages. Previously collected filtrate from Amicon spin column filter for pooled lavages for days 7, 21, 35, and 49 were resuspended in 25μL 10% sodium dodecyl sulfate solution with a protease inhibitor cocktail (Roche). Reduction of disulfides was performed by adding dithiothreitol (DTT) to the lavages to a final concentration of 20 mM and heating at 95° C for 10 min. After cooling to rt, alkylation of cysteines was performed by adding iodoacetamide to a final concentration of 40 mM and incubating in the dark for 30 min. Lavages were then centrifuged at 13000 g for 8 min to remove any undissolved matter. 12% aqueous phosphoric acid was added at 1:10 and mixed for a final concentration of 1.2% phosphoric acid. 165 μL of triethylammonium bicarbonate (TEAB) in 90:10 methanol:MilliQ water (S-Trap buffer) was added to lysed lavages and mixed. The mixture was then added to the S-Trap. The S-Trap was centrifuged at 2000 g for 30 sec to elute the S-Trap buffer and retain proteins in the S-Trap column. 150 μL of S-Trap buffer was added to the S-Trap and centrifuged at 2000 g for 30 sec; this process was repeated five times with the S-Trap column being rotated 180 degrees between repeated centrifugations. rLys-C (Promega) was added at 1:50 weight:weight to the S-Trap column, which was placed in a clean microcentrifuge tube. Digestion was performed by incubating at 37° C overnight. After digestion, the following solutions were added to the S-Trap in sequential order to elute peptides: 1) 40 μL 50mM TEAB in MilliQ water followed by centrifugation at 4000 g for 30 sec, 2) 40 μL 0.2% formic acid in MilliQ water followed by centrifugation at 4000 g for 30 sec, and 3) 35 μL of 60:40 acetonitrile:MilliQ water followed by centrifugation at 4000 g for one minute. Eluted peptide solutions were dried *in vacuo* at 60° C for 1 hr.

### Enzymatic Digestion of Murine Vaginal Lavages Using In-Solution Digestion

In-solution enzymatic digestion was also performed on a small aliquot of pooled murine vaginal lavages for days 35 and 49. Lavages were reduced and alkylated as outlined in the above S-Trap digestion protocol. Following reduction and alkylation, rLys-C (Promega) was added at 1:50 enzyme wt:sample wt to the lavage, which were then incubated at 37° C overnight and dried *in vacuo* at 60° C for 1 hr.

### Sample Desalting and Preparation for LC-MS/MS

Following enzymatic digestion, samples were resuspended in 100 μL 0.1% formic acid in MilliQ water. Samples were then desalted using C18 Resin ZipTip pipette tips (Millipore Sigma) per the manufacturer’s protocol, and desalted samples were dried *in vacuo* at 45° C for 30 min. After drying, samples were resuspended in 0.1% formic acid to a concentration of 0.5 μg/μL and centrifuged at 13000 g for 10 min to remove any particulates that may be present in the sample before being transferred to plastic LC-MS/MS vials. A 0.1% formic acid in MilliQ water solvent blank was also prepared for LC-MS/MS data acquisition.

### Data Acquisition via LC-MS/MS

Analysis was performed on a Q Exactive mass spectrometer (Thermo Fisher Scientific) coupled to an Agilent 1260 Infinity nanoLC system (Agilent Technologies). Samples were loaded onto an Acclaim Pepmap 100 C18 trap column (75 μm x 2 cm nanoViper, 3 μm, 100 Å) (Thermo Fisher Scientific) at 2 μL/min. After 10 min of washing with H_2_O w/ 0.1% formic acid (solvent A), separation was performed using a 60-min gradient at a flow rate of 0.25 μL/min on a Zorbax 300SB-C18 column (0.075 x 150 mm, 3.5 μm, 300 Å). The gradient was as follows: 60 min from 5 - 30% acetonitrile w/ 0.1% formic acid (solvent B), then 20 min from 30 - 60% B. The system was then maintained at 90% B for 10 min followed by a 15 min re-equilibration segment at 5% B. Data-dependent acquisition was used during the collection of mass spectra with a capillary temperature of 250° C and spray voltage of 1.5 kV. Full MS scans were collected at a mass resolution of 70,000 with a scan range of *m/z* 375 - 2000. Automatic gain control (AGC) target was set at 1e6 for a maximum injection time (IT) of 100 ms. The top 10 most intense peaks were selected for MS/MS analysis, with an isolation width of 1.5 *m/z*. MS/MS spectra were acquired at a resolution of 17,5000 with an AGC target of 1e5 and maximum IT of 50 ms. The first fixed mass was set at 100 *m/z*. Parent ions were fragmented at a normalized collision energy (NCE) of 27. Dynamic exclusion was set for 20 sec.

### Annotation of Microproteins Using MaxQuant

Protein annotation was performed on LC-MS/MS runs using the open-source platform MaxQuant (version 1.5.4.0) against UniProt proteomes for *Homo sapien, Mus muscula, and Rattus norvegicus* (release-01_2020). Default MaxQuant parameters were used unless otherwise specified, and Lys-C was specified as the digestion enzyme, allowing up to two missed cleavages. Label-free quantification was disabled due to insufficient replicates as a result of sample pooling. Carbamidomethylation was specified as a fixed modification, while oxidation and protein N-terminal acetylation were set as variable modifications, and a maximum of five modifications per peptide were allowed. The minimum peptide length allowed was set to seven, and the maximum peptide mass was 30 kDa. The minimum and maximum peptide length for unspecific search were set to 8 and 25, respectively. The peptide-spectrum match, protein, and site false discovery rates were all set to 1%.

### Retrospective Analysis of Vaginal Lavage Protein Profiles Acquired via MALDI-TOF MS

Following annotation in MaxQuant, results from the protein groups file (see MaxQuant documentation for output files) were filtered to remove any annotations with a mass greater than 32 kDa, less than two unique peptides, and any contaminants, which resulted in a total of 14 proteins (**Table 1**).

Protein profiles for the vaginal lavages were also manually inspected for each of the 14 proteins in R. A total of 1056 profiles were loaded using the MALDIquant R package for data preprocessing (five mice, weekly lavages collected from Day 0 - Day 56, 24 technical replicates per lavage; Day 56 lavage for Mouse 904 was absent). Parameters for the following preprocessing steps were left at their default values unless otherwise specified. All spectra were trimmed (MALDIquant::trim) to a range of 4000 - 20000 Da. Intensity transformation (MALDIquant::transformIntensity) was performed using the square root (‘sqrt’) method. Baseline smoothing (MALDIquant::smoothIntensity) was performed using the ‘SavitzkyGolay’ method. Baseline removal (MALDIquant::removeBaseline) was performed using the ‘TopHat’ method. Intensity normalization (MALDIquant::calibrateIntensity) was performed using the total ion current (‘TIC’) method. Peak detection (MALDIquant::detectPeaks) was performed using the minimum absolute deviation (‘MAD’) method of noise estimation and a signal to noise ratio of 3:1. Peak alignment (MALDIquant::alignPeaks) was performed with a signal to noise ratio of 3:1 and a tolerance of 0.2 Da; peaks were excluded if they were found to occur in less than 75% of the dataset. Preprocessing yielded a feature matrix containing 1939 features.

Six proteins annotated by MaxQuant could be found in this feature list as highlighted in **Table 1**. Of these six features, intensity over time for cystatin A and protein S100-A8 was plotted in **Figure 2a** and **Figure S1a**. Outliers were detected and removed using the interquartile range (IQR) method (**Equation 1**) prior to plotting. The regression of the mean intensity by day and mouse were also plotted using the ggplot2::geom_smooth function calculated with the ‘linear’ method (**Figure 2b** and **Figure S1b**).

### Validation of Cystatin A From Murine Tumors and Tissue Using Immunohistochemistry

Slides of tumors and reproductive tissues were subjected to heat-induced antigen retrieval after deparaffinization using sodium citrate at 100 °C for 30 min and inactivation of endogenous peroxidase activity using 0.3% H_2_O_2_/methanol for 15 min. After rinsing with phosphate buffered saline with Tween-20 (PBST), the slides were blocked with 5% horse serum (Vectastain ABC kit, Vector Laboratories, Inc.) diluted in 1% BSA/PBST at RT for 60 min. The tissue sections were incubated with cystatin A primary antibody (1:100, ThermoFisher PA5-75206) overnight at 4 °C. The next day, slides were rinsed with PBST prior to incubation with a biotinylated secondary antibody (Vectastain ABC kit, Vector Laboratories, Inc.) at 1:200 dilution in PBST for 60 min at rt. Slides were then rinsed and incubated in ABC solution (PBS: A: B=50:1:1) (Vectastain ABC kit, Vector Laboratories, Inc.) for 30 min at rt. For visualization of the immunoreactivity, all slides were subjected to chromogen 3’3-diaminobenzidine (DAB) (Vector Laboratories, Inc. Burlingame, CA) for 1 min and rinsed in running tap water for 10 min. Then slides were counterstained with hematoxylin and imaged using Nikon E600 Eclipse microscope with CMOS C-Mount microscope camera.

### Preparation of Cystatin A Antibody Functionalized Microtoroid

To specifically detect cystatin A in mouse samples, recombinant monoclonal cystatin A antibodies (Thermo Fisher Scientific, MA5-29200) were immobilized on the silica surface of the microtoroid through EDC/sulfo-NHS covalent coupling. Microtoroid optical resonators were fabricated as described previously using a combination of photolithography and etching steps^50^. The final structure was formed by melting the silica disk edge with a CO^2^ laser. Microtoroids were treated with 1% v/v APTES in 1 mM acetic acid for 10 min and incubated overnight with a solution of 100 mg/mL succinic anhydride in DMSO. After succinylation, the chip was rinsed with DMSO and ethanol before it was dried in a stream of nitrogen. EDC/sulfo-NHS 100 mM/100 mM in 0.1 M MES buffer, pH 5, was prepared freshly and immediately applied on the chip. After 10 min of EDC/NHS conjugation, the chip was washed with 10 mM sodium phosphate buffer (PBS), pH 7.4, and then incubated with cystatin A antibodies (50 μg/mL in PBS buffer) for 1 hr. The surface was subsequently quenched with 100 mM ethanolamine for 5 min to block residual amine-reactive groups. The antibody-coated chip was incubated in PBS buffer for further sensing experiments.

### Detection of Cystatin A Using FLOWER

Immediately prior to running FLOWER experiments, samples were thawed and aliquoted; 4-5 μL was removed from each sample and diluted a thousand fold for usage. The 1 μL sample was diluted 4000 fold. Remaining aliquots were flash-frozen in a cooling bath of dry ice/isopropyl acetone (BTC, 211315) and stored at −20 °C. FLOWER (**Figure 1c-f**) was used to investigate progressing cystatin A levels in murine vaginal lavage samples^16–18^. The antibody-immobilized toroid chip is mounted on an open microfluidic chamber which is designed to allow the passage of the optical fiber. Diluted murine samples are continuously perfused through the chamber at a steady flow rate of 100 μL/min. Here, the cystatin A antigen-antibody binding events are monitored in real time by adaptively tracking the resonance frequency of the microtoroid to obtain analyte binding curves. The monitoring of the shift in resonance wavelength of the microtoroid is recorded while steadily perfusing samples into the chamber. Each diluted murine sample is flowed for 5 min followed by 1 min of regeneration buffer (glycine-HCl, pH 3.0) to dissociate cystatin A from the antibodies and 5 min of sensing buffer (10 mM PBS, pH 7.4) rinsing. The recordings which last for more than 1 hr were segmented to extract the binding curves for murine samples from different timepoints. Extracted curves are further calibrated by subtracting the nearby buffer background. A representative curve of cystatin A binding to anti-cystatin A in PBS is shown in Fig. S4.

### Data Availability

MALDI-TOF MS protein profiles previously reported on by Galey *et al*. can be found at the following MassIVE repository: MSV000083628. Raw data from LC-MS/MS based bottom-up proteomics and parameters/results from MaxQuant can be found in the following MassIVE repository: MSV000088568. All code and data used for MALDI-TOF MS preprocessing and figure creation can be found in the following Github repository: https://github.com/gtluu/cystatin_a_figures.

## Acknowledgments

We would like to acknowledge Thu (Mi) Nguyen and Chandimal Pathmasiri for their expert input and assistance in performing LC-MS/MS based bottom-up proteomics experiments. JS acknowledges support from NIH R35GM137988. In addition, JS acknowledges support from Scialog^®^ and the Research Corporation for Science Advancement through an award sponsored by the Flinn Foundation. C.G. is supported by the Robert R. Shannon Graduate Student Endowed Scholarship in Optical Sciences. This publication was supported by the National Cancer Institute Award Number R01CA240423 (LMS and JEB) of the National Institutes of Health, the Research Corporation for Science Advancement Scialog^®^ Award #26222 (LMS). GTL was previously supported by NIH/NCCIH grant T32AT007533.

## References

1. Sung, H. et al. Global Cancer Statistics 2020: GLOBOCAN Estimates of Incidence and Mortality Worldwide for 36 Cancers in 185 Countries. CA: A Cancer Journal for Clinicians vol. 71 209–249 (2021).

2. Cancer of the Ovary - Cancer Stat Facts. https://seer.cancer.gov/statfacts/html/ovary.html.

3. Cancer of the Endometrium - Cancer Stat Facts. https://seer.cancer.gov/statfacts/html/corp.html.

4. Common Cancer Sites - Cancer Stat Facts. https://seer.cancer.gov/statfacts/html/common.html.

5. Luu, G. T. & Sanchez, L. M. Toward improvement of screening through mass spectrometry-based proteomics: ovarian cancer as a case study. Int. J. Mass Spectrom. 469, (2021).

6. Swiatly, A., Plewa, S., Matysiak, J. & Kokot, Z. J. Mass spectrometry-based proteomics techniques and their application in ovarian cancer research. J. Ovarian Res. 11, 1–13 (2018).

7. Mukherjee, S., Sundfeldt, K., Borrebaeck, C. A. K. & Jakobsson, M. E. Comprehending the Proteomic Landscape of Ovarian Cancer: A Road to the Discovery of Disease Biomarkers. Proteomes 9, 25 (2021).

8. Elzek, M. A. & Rodland, K. D. Proteomics of ovarian cancer: functional insights and clinical applications. Cancer Metastasis Rev. 34, 83–96 (2015).

9. Brinton, L. T., Brentnall, T. A., Smith, J. A. & Kelly, K. A. Metastatic Biomarker Discovery Through Proteomics. Cancer Genomics Proteomics 9, 345–355 (2012).

10. Costas, L. et al. New perspectives on screening and early detection of endometrial cancer.Int. J. Cancer 145, 3194–3206 (2019).

11. Galey, M. M. et al. Detection of Ovarian Cancer Using Samples Sourced from the Vaginal Microenvironment. J. Proteome Res. 19, 503–510 (2020).

12. Petukhova, V. Z. et al. Whole Cell MALDI Fingerprinting Is a Robust Tool for Differential Profiling of Two-Component Mammalian Cell Mixtures. J. Am. Soc. Mass Spectrom. 30, 344–354 (2019).

13. Lisio, M.-A., Fu, L., Goyeneche, A., Gao, Z.-H. & Telleria, C. High-Grade Serous Ovarian Cancer: Basic Sciences, Clinical and Therapeutic Standpoints. Int. J. Mol. Sci. 20, (2019).

14. Li, C., Chen, L., McLeod, E. & Su, J. Dark mode plasmonic optical microcavity biochemical sensor. Photonics Research vol. 7 939 (2019).

15. Su, J. Label-free Single Molecule Detection Using Microtoroid Optical Resonators. Journal of Visualized Experiments (2015) doi:10.3791/53180.

16. Su, J., Goldberg, A. F. G. & Stoltz, B. M. Label-free detection of single nanoparticles and biological molecules using microtoroid optical resonators. Light: Science & Applications 5, e16001–e16001 (2016).

17. Su, J. Label-free single exosome detection using frequency-locked microtoroid optical resonators. ACS Photonics 2, 1241–1245 (2015).

18. Ozgur, E. et al. Ultrasensitive detection of human chorionic gonadotropin using frequency locked microtoroid optical resonators. Anal. Chem. 91, 11872–11878 (2019).

19. Hao, S. & Su, J. Noise-Induced Limits of Detection in Frequency Locked Optical Microcavities. Journal of Lightwave Technology vol. 38 6393–6401 (2020).

20. Gibb, S. & Strimmer, K. MALDIquant: a versatile R package for the analysis of mass spectrometry data. Bioinformatics 28, 2270–2271 (2012).

21. Panich, S. et al. Label-Free Pb(II) Whispering Gallery Mode Sensing Using Self-Assembled Glutathione-Modified Gold Nanoparticles on an Optical Microcavity. Analytical Chemistry vol. 86 6299–6306 (2014).

22. Suebka, S., Nguyen, P.-D., Gin, A. & Su, J. How Fast It Can Stick: Visualizing Flow Delivery to Microtoroid Biosensors. ACS Sens 6, 2700–2708 (2021).

23. Wilson, D. A. et al. Multicenter Evaluation of the Bruker MALDI Biotyper CA System for the Identification of Clinically Important Bacteria and Yeasts. Am. J. Clin. Pathol. 147, 623–631 (2017).

24. Milani, P. et al. The utility of MASS-FIX to detect and monitor monoclonal proteins in the clinic. Am. J. Hematol. 92, 772–779 (2017).

25. Kohlhagen, M. et al. Automation and validation of a MALDI-TOF MS (Mass-Fix) replacement of immunofixation electrophoresis in the clinical lab. Clin. Chem. Lab. Med. 59, 155–163 (2020).

26. Murray, D. L. et al. Mass spectrometry for the evaluation of monoclonal proteins in multiple myeloma and related disorders: an International Myeloma Working Group Mass Spectrometry Committee Report. Blood Cancer J. 11, 24 (2021).

27. Mellors, P. et al. MASS-FIX for the diagnosis of plasma cell disorders: A single institution experience of 4118 patients. Blood 136, 48–49 (2020).

28. Mellors, P. W. et al. MASS-FIX for the detection of monoclonal proteins and light chain N-glycosylation in routine clinical practice: a cross-sectional study of 6315 patients. Blood Cancer J. 11, 50 (2021).

29. Myeloma - Cancer Stat Facts. https://seer.cancer.gov/statfacts/html/mulmy.html.

30. Kazandjian, D. Multiple myeloma epidemiology and survival: A unique malignancy. Semin. Oncol. 43, 676–681 (2016).

31. Rinne, A. Epidermal SH-protease inhibitor (ACPI, cystatin A) in cancer. A short historical review. Pathol. Res. Pract. 206, 259–262 (2010).

32. Machleidt, W. et al. Protein inhibitors of cysteine proteinases. II. Primary structure of stefin, a cytosolic protein inhibitor of cysteine proteinases from human polymorphonuclear granulocytes. Hoppe Seylers Z. Physiol. Chem. 364, 1481–1486 (1983).

33. Yamazaki, M., Ishidoh, K., Kominami, E. & Ogawa, H. Genomic structure of human cystatin A. DNA Seq. 8, 71–76 (1997).

34. Martin, J. R. et al. The three-dimensional solution structure of human stefin A. J. Mol. Biol. 246, 331–343 (1995).

35. Räsänen, O., Järvinen, M. & Rinne, A. Localization of the human SH-protease inhibitor in the epidermis. Immunofluorescent studies. Acta Histochem. 63, 193–196 (1978).

36. Hsieh, W. T., Fong, D., Sloane, B. F., Golembieski, W. & Smith, D. I. Mapping of the gene for human cysteine proteinase inhibitor stefin A, STF1, to chromosome 3cen-q21. Genomics 9, 207–209 (1991).

37. Kos, J. et al. Cysteine Proteinase Inhibitors Stefin A, Stefin B, and Cystatin C in Sera from Patients with Colorectal Cancer: Relation to Prognosis. Clin. Cancer Res. 6, 505–511 (2000).

38. Parker, B. S. et al. Primary tumour expression of the cysteine cathepsin inhibitor Stefin A inhibits distant metastasis in breast cancer. J. Pathol. 214, 337–346 (2008).

39. Lah, T. T. et al. The Expression of Lysosomal Proteinases and Their Inhibitors in Breast Cancer: Possible Relationship to Prognosis of the Disease. Pathol. Oncol. Res. 3, 89–99 (1997).

40. Kuopio, T. et al. Cysteine proteinase inhibitor cystatin A in breast cancer. Cancer Res. 58, 432–436 (1998).

41. Leinonen, T. et al. Biological and prognostic role of acid cysteine proteinase inhibitor (ACPI,cystatin A) in non-small-cell lung cancer. J. Clin. Pathol. 60, 515–519 (2007).

42. Vasilopoulos, Y. et al. Association analysis of the skin barrier gene cystatin A at the PSORS5 locus in psoriatic patients: evidence for interaction between PSORS1 and PSORS5. Eur. J. Hum. Genet. 16, 1002–1009 (2008).

43. Kennedy, K. D., AnithaChristy, S. A., Buie, L. K. & Borrás, T. Cystatin a, a potential common link for mutant myocilin causative glaucoma. PLoS One 7, e36301 (2012).

44. Abrahamson, M., Barrett, A. J., Salvesen, G. & Grubb, A. Isolation of six cysteine proteinase inhibitors from human urine. Their physicochemical and enzyme kinetic properties and concentrations in biological fluids. J. Biol. Chem. 261, 11282–11289 (1986).

45. Abrahamson, M., Alvarez-Fernandez, M. & Nathanson, C.-M. Cystatins. Biochem. Soc. Symp. 179–199 (2003).

46. Lah, T. T. et al. Cystatins and stefins in ascites fluid from ovarian carcinoma. Cancer Lett. 61, 243–253 (1992).

47. Kastelic, L. et al. Stefin B, the major low molecular weight inhibitor in ovarian carcinoma. Cancer Lett. 82, 81–88 (1994).

48. National Cancer Institute. SEER Cancer Stat Facts: Ovarian Cancer. https://seer.cancer.gov/statfacts/html/ovary.html.

49. Diz, A. P., Truebano, M. & Skibinski, D. O. F. The consequences of sample pooling in proteomics: an empirical study. Electrophoresis 30, 2967–2975 (2009).

50. Armani, D. K., Kippenberg, T. J., Spillane, S. M. & Vahala, K. J. Ultra-high-Q toroid microcavity on a chip. Nature 421, 925–928 (2003).

