## Supplemental Information for "An integrated approach to protein discovery and detection from complex biofluids"

Figure S1: Intensity and linear regression of mean intensity for protein S100-A8 over time.……S2

Figure S2: Increase in tumor burden in mice over time……………………………………………S3

Figure S3: FLOWER relative shifts over time………………………………………………………S4

Figure S4: Detection of known concentrations of cystatin A via FLOWER…………………………S6

**Supplemental Information**

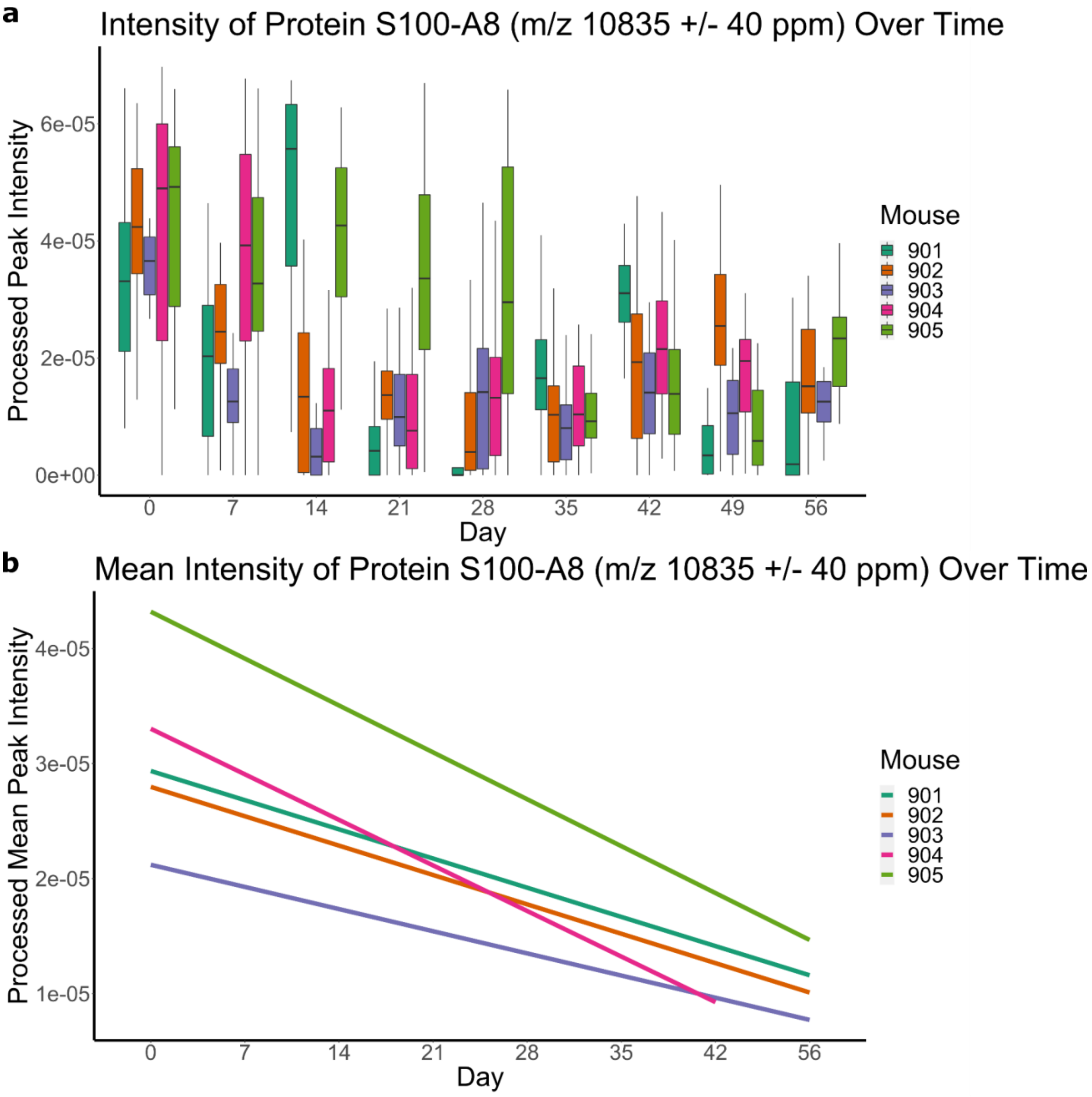


**Figure S1**: (a) Box plot showing the intensity of protein S100-A8 at each corresponding time point. (b) Linear regression trend lines for the mean intensities of protein S100-A8 at each time point showing downregulation.

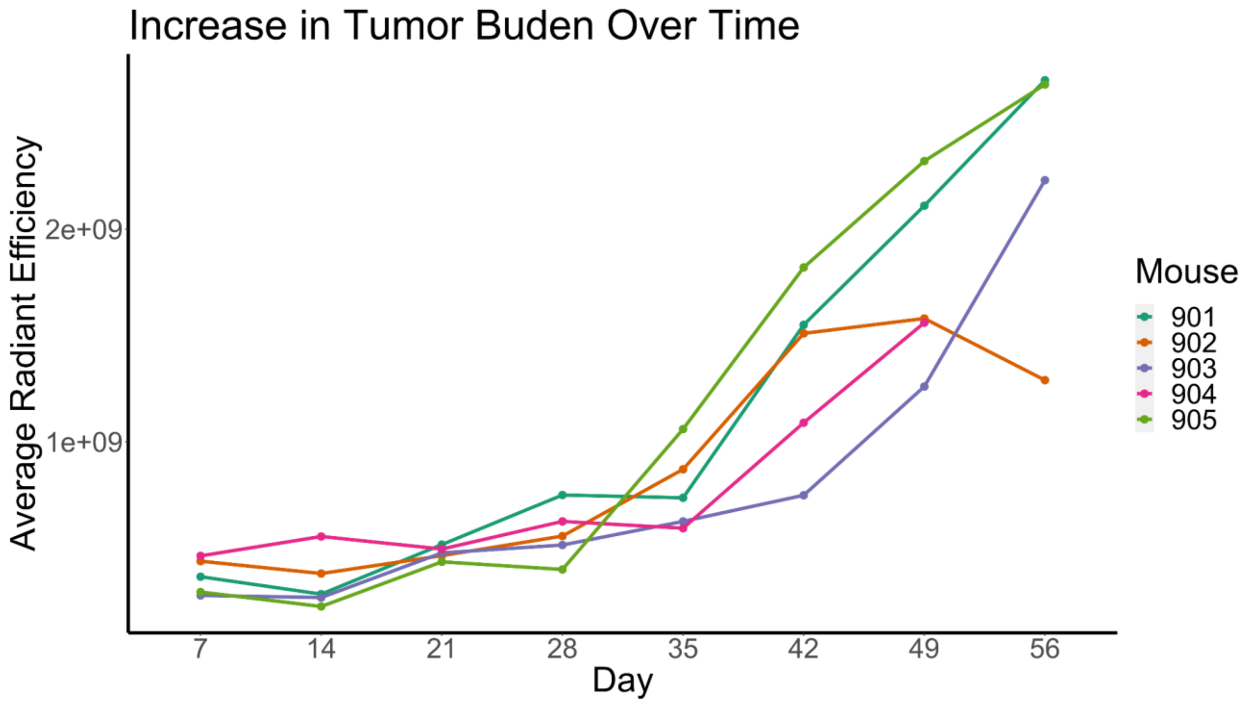


**Figure S2**: Tumor burden in five biological replicate xenograft murine models transfected with OVCAR-8-RFP tumors. Tumor burden was measured weekly over 56 days by IVIS.

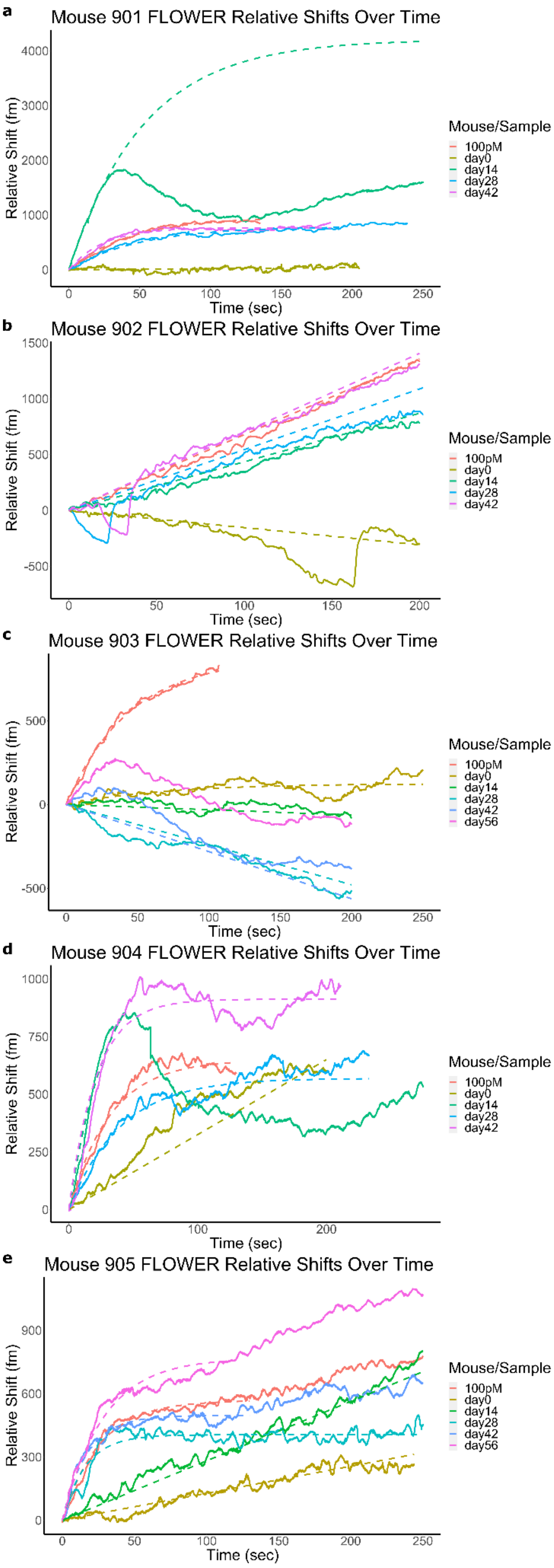
a
Mouse 901 FLOWER Relative Shifts Over Time
Relative Shift (fm)
4000
3000
2000
1000
0
0
50
100
150
200
250
Time (sec)
Mouse/Sample
100pM
day0
day14
day28
day42
b
Mouse 902 FLOWER Relative Shifts Over Time
Relative Shift (fm)
1500
1000
500
0
-500
0
50
100
150
200
Time (sec)
Mouse/Sample
100pM
day0
day14
day28
day42
c
Mouse 903 FLOWER Relative Shifts Over Time
Relative Shift (fm)
500
0
-500
0
50
100
150
200
250
Time (sec)
Mouse/Sample
100pM
day0
day14
day28
day42
day56
d
Mouse 904 FLOWER Relative Shifts Over Time
Relative Shift (fm)
1000
750
500
250
0
0
100
200
Time (sec)
Mouse/Sample
100pM
day0
day14
day28
day42
e
Mouse 905 FLOWER Relative Shifts Over Time
Relative Shift (fm)
900
600
300
0
0
50
100
150
200
250
Time (sec)
Mouse/Sample
100pM
day0
day14
day28
day42
day56

**Figure S3**: Label-free detection of cystatin A using a microtoroid resonator. (a-e) Cystatin A binding curves for Mouse 901-905. Similar to MS protein profiles, Mouse 903 displayed a strange lack of detectable cystatin A. Several time points (i.e Mouse 901 Day 14 and Mouse 904 Day 14) also appeared to have abnormal binding curves, most likely owed to biological variability and/or murine model behavior during collection of murine vaginal lavages. The dashed curves represent fits to the experimental data using Equation (2) unless the r-squared value with a fit to Equation (1) was better in which case Equation (1) was used, which has fewer fitting parameters than Equation (2). In both cases, the fit was used to obtain the initial slope of the curve.

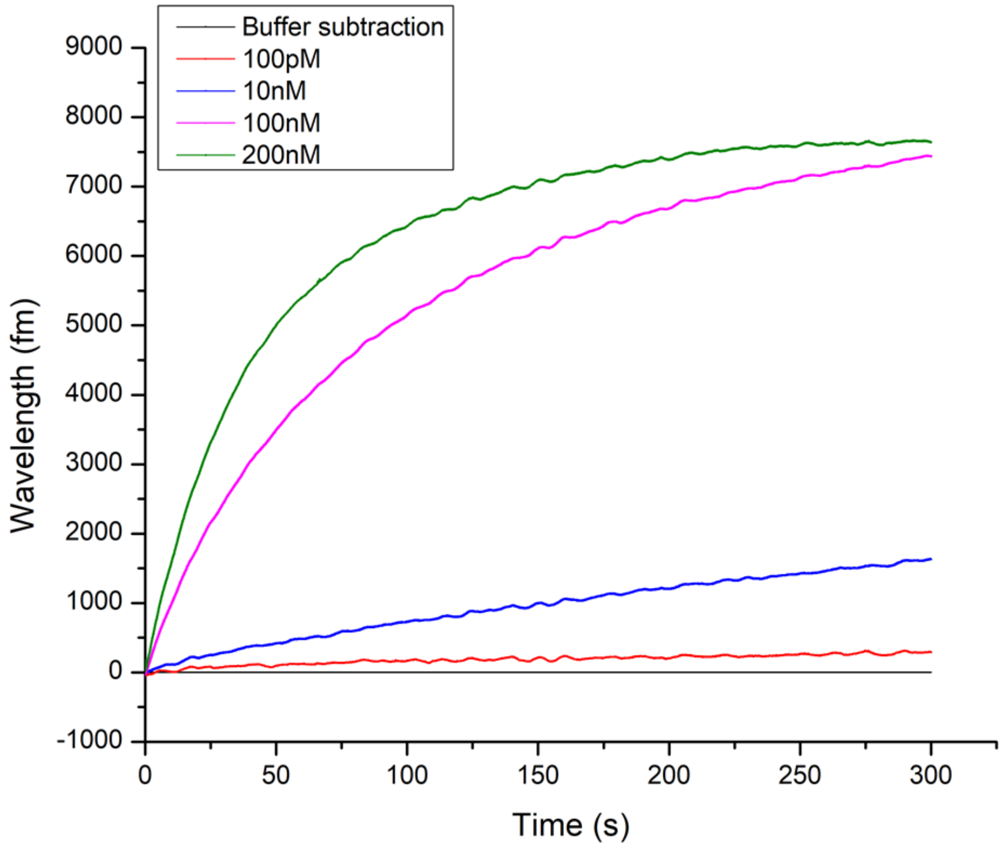


**Figure S4:** Detection of known concentrations of cystatin-A binding to anti-cystatin-A using FLOWER. Experiments were performed in PBS.
